# A precision-cut lung slice platform for evaluating respiratory virus replication dynamics

**DOI:** 10.64898/2026.05.28.728430

**Authors:** Jovanna A. Fusco, Min Liu, Devra Huey, Emily M. King, Amanda R. Panfil, Kara N. Corps, Ian C. Davis, Andrew S. Bowman, Cody J. Warren

## Abstract

Understanding respiratory virus replication, tropism, and disease mechanisms requires experimental systems that balance physiological relevance with scalability. Although immortalized cell lines remain widely used, they fail to capture the cellular complexity of the lung, while animal models—though informative—are often costly, low-throughput, and limited in their ability to model human disease. Here, we describe a rapid and scalable precision-cut lung slice (PCLS) platform that overcomes many of these limitations. Our workflow generates infection-ready lung tissue slices within 24 hours and maintains tissue viability over extended culture periods. Using influenza A virus as a model pathogen, we demonstrate that PCLS support robust viral replication, recapitulate characteristic infection-associated pathology, and elicit localized immune cell responses within the respiratory epithelium. Together, these features establish PCLS as a versatile and physiologically relevant platform for studying the biology of influenza virus and other respiratory pathogens.

## Introduction

Respiratory viruses remain a major global cause of morbidity and mortality.^1^ Influenza viruses alone are responsible for substantial seasonal disease burden and periodically give rise to pandemics with profound and lasting global health consequences. Despite decades of study, these viruses continue to surprise us, as continued viral evolution, reassortment, and cross-species transmission generate novel viruses with unique molecular properties and disease characteristics. The rapid worldwide spread of clade 2.3.4.4b H5N1 influenza viruses, coupled with increased mammalian susceptibility, is a perfect case-in-point.^2,3^

As influenza and other respiratory viruses continue to exert staggering global health burdens, new technologies to evaluate virus biology become critical.^4^ Progress in developing effective therapies has been hindered by the lack of physiologically relevant model systems that recapitulate the structural and cellular complexity of the lung. Lung function depends on the coordinated activity of many specialized cell types that maintain structure, enable gas exchange, and defend against infection. Conventional two-dimensional culture models fail to capture this complexity, limiting their translational value, while animal models, though informative—are costly, low-throughput, and may not accurately recapitulate human respiratory biology or disease outcomes.

Precision-cut lung slices (PCLS) provide a physiologically relevant *ex vivo* platform that bridges the gap between traditional cell culture and *in vivo* models.^5^ Unlike monolayer cultures, PCLS preserve the native three-dimensional architecture of the lung, including intact airways, alveolar structures, and the surrounding extracellular matrix. These slices retain diverse resident cell types—epithelial, stromal, vascular, and smooth muscle—as well as functional immune cells such as macrophages^6^ and neutrophils^7^ that contribute to early host responses. By maintaining this complex tissue microenvironment, PCLS enable disease modeling under conditions that closely reflect the intact lung.

A key advantage of the PCLS approach is its efficiency and scalability, enabling the generation of multiple uniform tissue slices from a single lung to support high-throughput, reproducible experimentation. The versatility of this 3D model further extends its value—it can be established from both animal and human lungs, including tissues derived from healthy donors or individuals with underlying disease, thereby enabling direct comparison across species and disease states.^8^

Here, we describe a standardized pipeline for the high-throughput generation of precision-cut lung slices. Culture conditions were optimized to maintain tissue viability for extended periods, and tissues recovered from cryopreservation retained viability—enabling opportunities for PCLS biobanking and resource sharing. Using influenza A virus (IAV) as a proof-of-principle model, we demonstrate that PCLS provides a robust platform to study respiratory virus replication, host responses, and infection-associated tissue pathology. Together, these advances establish a versatile, reproducible system for respiratory virus research that combines physiological relevance with experimental scalability.

## Results

### Design and optimization of a precision-cut lung slice pipeline for respiratory virus replication studies

Respiratory viruses are often studied in immortalized or primary airway epithelial cells, but these models lack the structural complexity of intact lung tissue. Precision-cut lung slices (PCLS) offer a physiologically relevant *ex vivo* platform that preserves native tissue architecture.^9,10^ To create a scalable and reproducible system, we developed and validated a PCLS pipeline using IAV as a model respiratory pathogen, with standardized readouts to assess viral replication dynamics and host responses (**Fig. 1A**).

**Figure 1.**
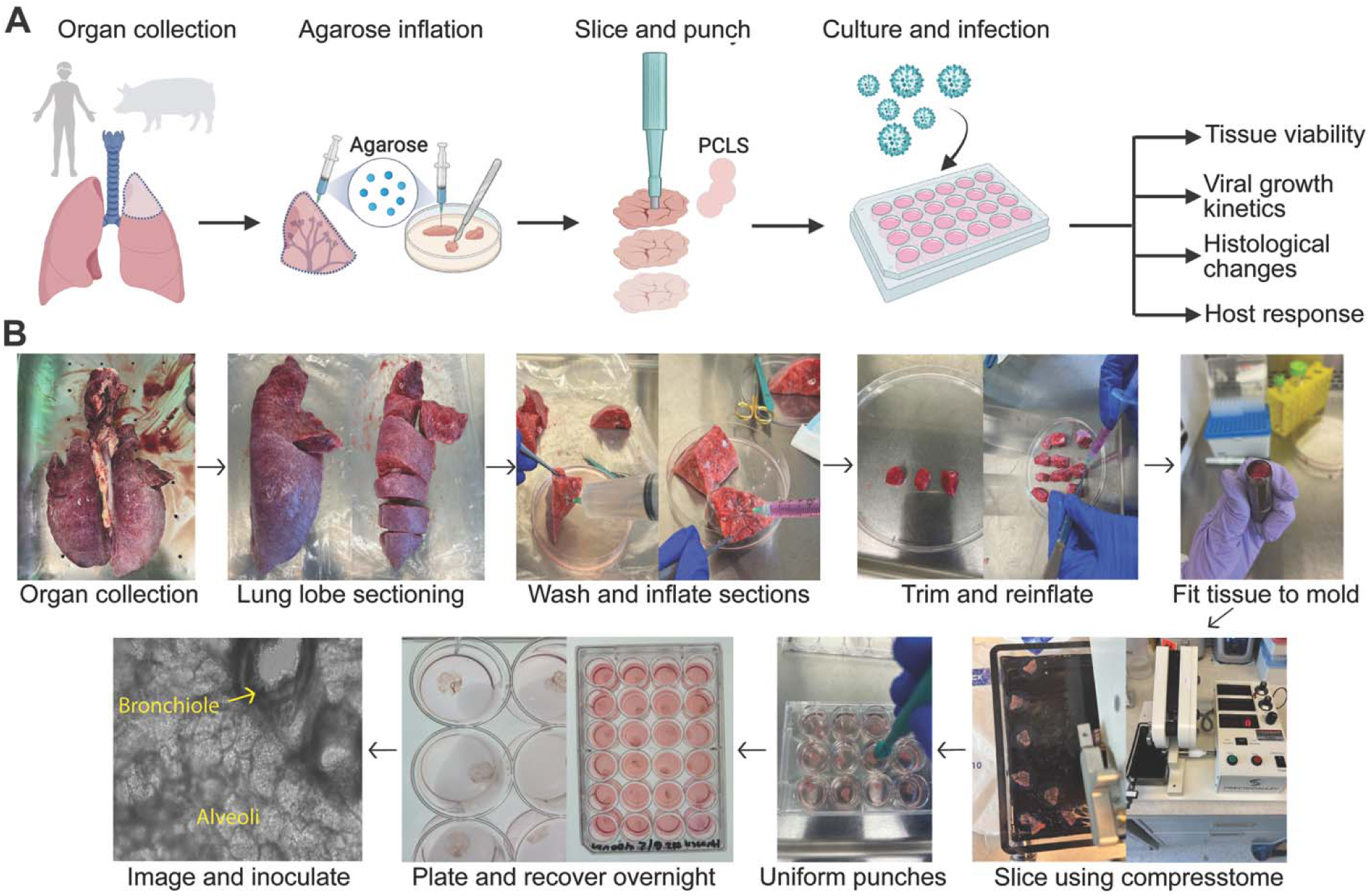
Design and optimization of a precision-cut lung slice pipeline for respiratory virus replication studies. (**A**) Schematic overview of the precision-cut lung slice workflow and experimental readouts. (**B**) Opportunistically collected lungs are sectioned, perfused with PBS through the airways, and inflated with low-melting-point agarose to stabilize the tissue. After trimming and reinflation, samples are embedded in a specimen tube mold and sectioned using a Compresstome®. Generated slices are standardized with a coring tool to produce uniform punches, which are plated in 24-well culture dishes and allowed to recover overnight. The resulting lung slices are then ready for downstream applications. Panels include representative images of each step.

The PCLS pipeline begins with opportunistic sampling of donor lungs, which can be obtained from research organ donations, explanted transplant tissue, or animal sources. For this study, we used swine lungs due to their accessibility and close similarity to human lung size and architecture.^11,12^ Swine lungs were opportunistically collected from a meat processing facility at harvest and transported to the laboratory, where lobes were isolated and prepared for slicing. Perfusion with PBS cleared residual blood and airway debris, while inflation with low-melting-point agarose stabilized alveolar structures and facilitated uniform sectioning. Stabilized tissues were then trimmed, re-inflated, and embedded in cylindrical molds for slicing using a Compresstome®, although other vibrating microtomes yielded comparable results. This approach reproducibly generated slices of consistent thickness (400 μm). To further standardize tissue dimensions across experiments, slices were cored with an 8 mm punch biopsy, which enabled consistent sizing and uniform handling in multi-well culture formats. The resulting PCLS recovered overnight in culture and were infection-ready within 24 hours of initial tissue harvest (**Fig. 1B**).

### Precision-cut lung slices support tissue viability and respiratory virus replication

To evaluate the stability of PCLS in culture, we first assessed tissue health using a luminescence-based assay that detects lactate dehydrogenase (LDH) release into the medium. Because LDH is retained in the cytoplasm of intact cells, its extracellular accumulation provides a quantitative measure of cell damage. Across a 5-day culture window, LDH release from both fresh and cryopreserved PCLS remained low and comparable to baseline levels, in contrast to a positive control (+; purified LDH enzyme) that established the assay’s upper detection limit (**Fig. 2A**). These results confirm that under standardized culture conditions, PCLS (1) remain viable for extended periods, (2) are amenable to cryopreservation and intact upon thaw, and (3) are suitable for downstream applications within a relatively broad timeframe.

**Figure 2.**
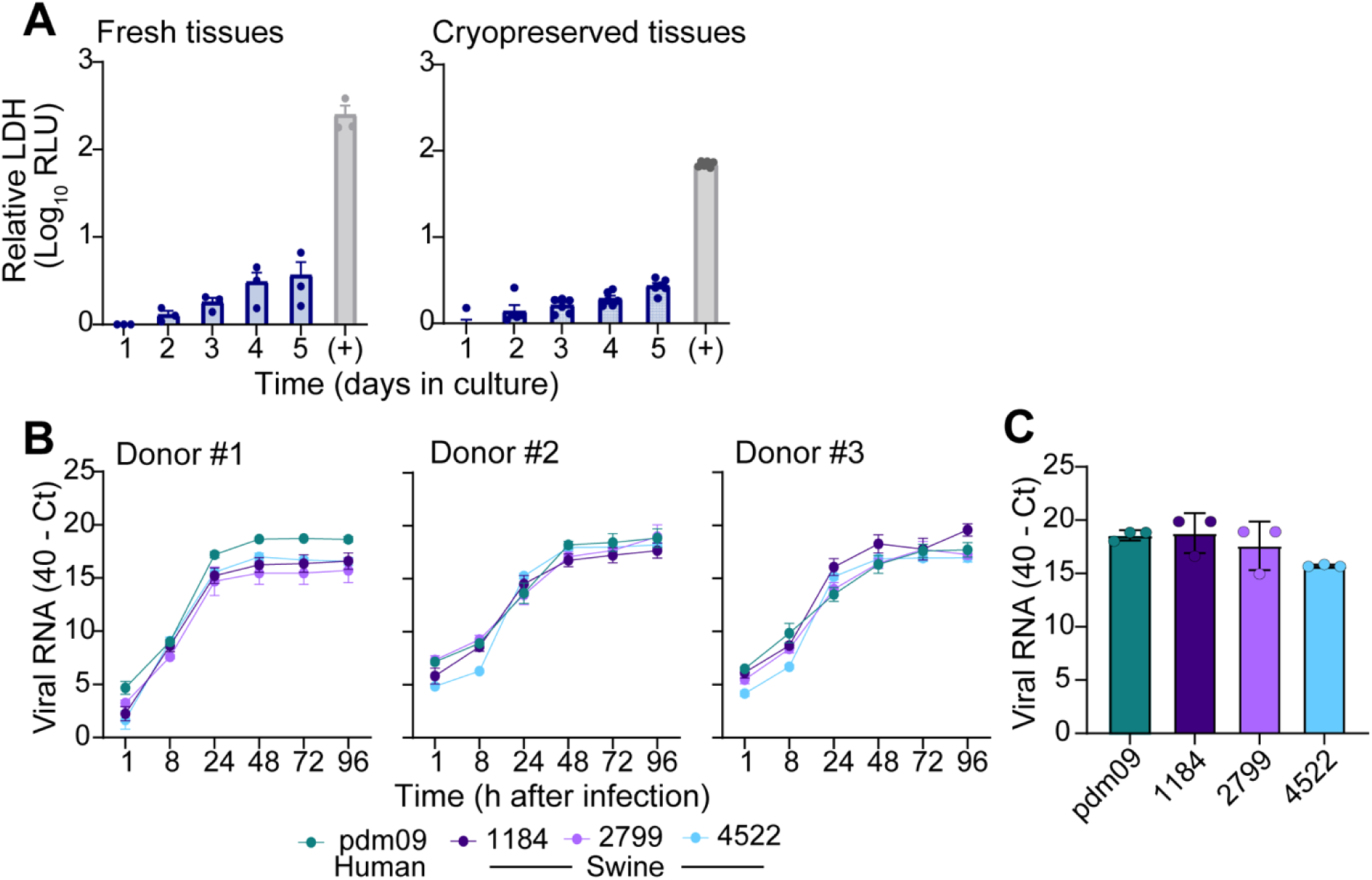
Precision-cut lung slices support tissue viability and respiratory virus replication. Opportunistically collected lungs from otherwise healthy swine sampled at slaughter were processed for precision-cut lung slices (PCLS). (**A**) Lactate dehydrogenase (LDH) assays were performed daily to assess tissue viability over time in culture. Purified LDH enzyme was included as a positive control to demonstrate the upper limit of the assay. For fresh tissues, error bars represent the mean +/- SEM from three independent donors, each with three replicates per donor. For cryopreserved tissues, error bars represent the mean +/- SEM from one donor with six replicates. (**B**) Swine PCLS were exposed to IAV isolated from humans (pdm09) or swine (1184, 2799, 4522) at equivalent quantities (3x10^4^ plaque forming units [PFU]). Culture media were collected at the indicated time points, and cell-free viral RNA levels were estimated by rRT-PCR. Data show the mean +/- SEM from three independent PCLS per donor tissue. (**C**) At 96 hours post-infection, three PCLS per donor were pooled, homogenized, and total RNA was extracted to quantify cell-associated viral RNA. Data are presented as mean ± SEM from three donor tissues.

Having established tissue viability, we next verified that swine PCLS could support infection with diverse IAV isolates from both human and endemic swine lineages. Twenty-four hours after harvest, tissues from three independent donors were inoculated with equivalent viral doses, washed extensively, and monitored for viral replication over time by quantifying viral RNA in culture supernatants using real-time reverse transcriptase PCR (rRT-PCR). All endemic swine viruses replicated robustly in PCLS, showing steadily increasing viral RNA levels throughout the time-course. Replication kinetics of the 2009 H1N1 pandemic human isolate (“pdm09”) mirrored those of swine viruses (**Fig. 2B**), consistent with expectations given its swine origin.^13–15^ To determine whether these findings reflected intracellular viral RNA accumulation levels, we homogenized PCLS at 96 hours post inoculation and measured intracellular viral RNA. Viral RNA accumulation in tissue lysates recapitulated supernatant data, with no remarkable differences observed between swine- and human-derived IAV isolates (**Fig. 2C**). Collectively, these results demonstrate that swine PCLS robustly support replication of both endemic swine and human IAVs, validating the system for comparative assessments of tissue level virus susceptibility.

### Precision-cut lung slices recapitulate infection-associated tissue damage and immune cell infiltration

To determine whether PCLS capture histopathological features of viral infection, we performed routine hematoxylin and eosin (H&E) staining on tissues following mock treatment or IAV inoculation. All sections were evaluated by a board-certified veterinary pathologist specializing in pulmonary infectious disease.

Mock-exposed PCLS maintained expected pulmonary architecture, including intact bronchioles and associated blood vessels, alveoli, and interstitium, as well as cellular populations such as airway epithelial cells, pneumocytes, alveolar macrophages, and normal lymphocytes found adjacent to airways (bronchiole-associated lymphoid tissue, BALT). Day 3 mock tissues retained these features with appropriate cytosolic and nuclear characteristics expected in normal, viable pulmonary cell populations (“Mock” in **Fig. 3A**). Together with preserved metabolic activity (**Fig. 2A**), these findings indicate that PCLS maintain both cellular viability and structural integrity over several days in culture.

**Figure 3.**
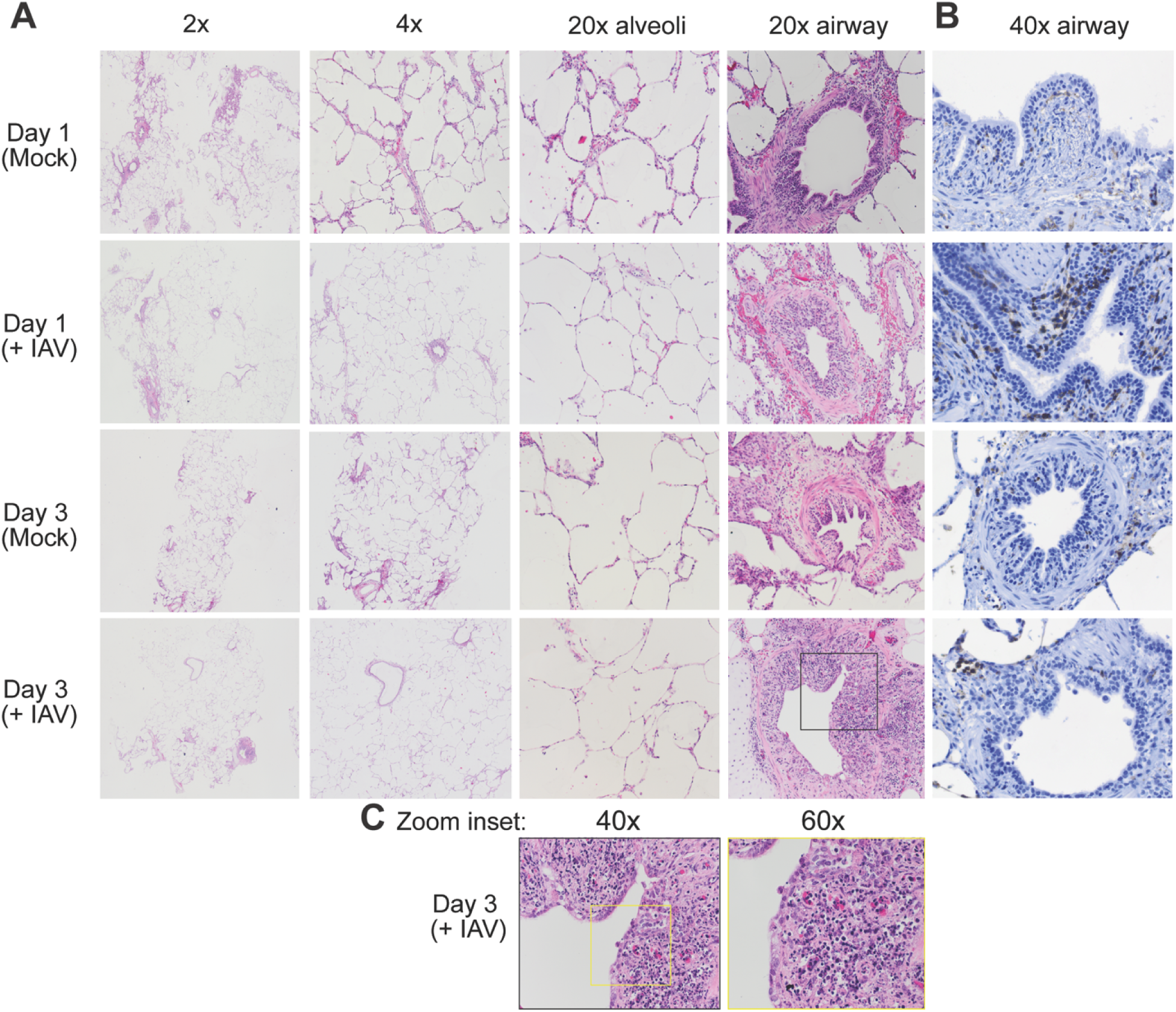
Precision-cut lung slices recapitulate infection-associated tissue damage and immune cell infiltration. (**A**) Representative hematoxylin and eosin (H&E) staining of swine PCLS at 1- or 3-days post-infection with influenza A virus (A/Puerto Rico/8/34, “PR8”; 3x10^4^ PFU) or mock treatment. Images are shown at low to high magnification (2x, 4x, 20x), highlighting alveolar regions and large airways containing bronchioles and bronchi. (**B**) Labeling of CD45+ immune cells demonstrates moderate inflammatory infiltrate in bronchiolar mucosa and adjacent interstitium on Day 1 postinfection, followed by severe infiltration and obscuring of the mucosa and adjacent tissues on Day 3. Inflammatory cells are mixed with necrotic debris from death of bronchiolar epithelial cells. Large numbers of IAV+ cells are present in affected tissue compartments. (**C**) High-magnification insets (40x and 60x) of bronchiolar epithelium demonstrate epithelial necrosis and damage and inflammatory cellular infiltration following infection.

In contrast, IAV-infected PCLS exhibited clear inflammatory and structural alterations. By day 1, infected tissues showed infiltration of CD45+ inflammatory cells crossing the bronchiolar epithelium and lamina propria and extending into the adjacent peri-bronchiolar interstitium, resulting in thickening of adjacent alveolar septa and focal disruption of bronchiolar mucosa (“+ IAV” in **Fig. 3A** and **3B**). By Day 3, lesions had progressed, with near-complete obscuring and/or loss of bronchiolar epithelium by inflammatory cells and marked attenuation of remaining pseudostratified columnar epithelial cells into a flattened, squamous morphology. The distinction between the bronchiolar mucosa and surrounding tissues is blurred by inflammation, edema, and necrotic cellular and nuclear debris (**Fig. 3A-C**). Together, these observations demonstrate that PCLS recapitulate the hallmark inflammatory and structural changes induced following acute IAV infection.

## Discussion

Advanced lung models have substantially improved our ability to study respiratory virus infection and host response. For example, primary airway epithelial cultures grown at the air–liquid interface (ALI) differentiate into a stratified epithelium that closely resembles the human respiratory tract.^16,17^ These physiologically relevant systems have been used to investigate a range of respiratory pathogens, including severe acute respiratory syndrome coronavirus 2 (SARS-CoV-2),^18^ respiratory syncytial virus (RSV),^19^ human rhinoviruses,^20^ and influenza A and influenza D viruses.^21,22^ Despite their utility, ALI cultures have several limitations: (1) extended culture times (≈4 weeks) are required to achieve full differentiation, (2) they primarily contain epithelial cells and lack interactions with stromal or vascular compartments, and (3) they do not include resident immune cells that play key roles in respiratory virus pathogenesis. These limitations are largely overcome by PCLS, which preserve the native lung architecture and cellular diversity (reviewed in ^23^). PCLS are infection-ready within 24 hours of tissue collection (**Fig. 1** and **Fig. 2B**) and phenocopy the cellular complexity of intact lung tissue (**Fig. 3**), enabling modeling of tissue injury and host response under physiologically relevant conditions. Furthermore, hundreds of uniform slices can be generated from a single lung, cryopreserved (**Fig. 2A**), and used on demand—addressing logistical challenges in tissue access and experimental reproducibility.

We show that swine PCLS support robust replication of both endemic swine and human IAV strains (**Fig. 2B–C**). Histological analyses further revealed infiltration of CD45+ inflammatory cells in the pulmonary interstitium and bronchiolar mucosal tissues, and a mixed inflammatory cell population consistent with neutrophils, macrophages, and lymphocytes, confirming that this system recapitulates localized, tissue-level immune responses to IAV infection (**Fig. 3**). These findings emphasize PCLS as a powerful ex vivo system for investigating influenza virus biology and pathogenesis. Consistent with this utility, our group recently demonstrated robust replication of bovine- and swine-origin influenza D virus isolates in human PCLS tissue, further supporting PCLS as a relevant platform for evaluating the replication capacity and zoonotic potential of emerging influenza viruses.^22^

Swine represent a critical animal model given their susceptibility to both avian- and mammalian-origin IAV and their recognized role in virus evolution.^24^ As the United States continues to confront widespread H5N1 outbreaks in domestic livestock,^3^ the swine PCLS model described here provides a tractable platform to compare susceptibility among avian- and mammalian-derived isolates and assess viral adaptation potential at the tissue level. The scalability and reproducibility of this approach enable experimental insights to be obtained rapidly, without reliance on resource-intensive animal studies. Moreover, extending this system to human-derived PCLS offers opportunities for therapeutic screening, as has been demonstrated for SARS-CoV-2.^25–27^

Together, this work establishes a standardized and scalable PCLS platform that enables physiologically relevant investigation of influenza virus infection and host response in intact lung tissue. Beyond its immediate applications in influenza virus research, this framework can be readily adapted to study diverse respiratory pathogens and evaluate therapeutic interventions. By enhancing the accessibility and reproducibility of PCLS, this study provides a foundation for physiologically relevant lung models that advance discovery in influenza virus biology, cross-species transmission, and disease pathogenesis While PCLS provide a powerful and physiologically relevant platform for studying influenza virus infection in intact lung tissue, several considerations merit discussion. In the current study, slices were maintained under submerged culture conditions, which do not fully recapitulate the air–liquid interface of the respiratory tract. Future adaptations incorporating transwell-based culture systems may better model airway-exposed infections, as demonstrated in other 3D respiratory systems.^28^ Additionally, because PCLS are derived from excised lung tissue, the immune compartment is limited to resident cells present at the time of collection. Although localized immune activation and inflammatory cell responses were observed following influenza virus infection (**Fig. 3**), this system does not capture immune cell recruitment from distal sites. Co-culture strategies pairing autologous PCLS with donor-matched immune populations represent a promising avenue to address this limitation.^29^ Finally, access to high-quality lung tissue can constrain experimental throughput; however, our demonstration that cryopreservation preserves tissue viability (**Fig. 2A**) establishes a practical path toward PCLS biobanking and on-demand use. Collectively, these considerations highlight opportunities for continued refinement while underscoring the utility of PCLS as a flexible and informative platform for influenza virus research.

## Resource availability

### Lead contact

Requests for further information and resources should be directed to and will be fulfilled by the lead contact, Cody Warren.

### Materials availability

Key resources generated in this study are available from the lead contact with a completed materials transfer agreement.

### Data availability

Any additional information required to reanalyze the data reported in this study is available from the lead contact upon request.

## Acknowledgments

This work was supported by the United States Department of Agriculture (USDA) National Institute of Food and Agriculture (NIFA) under contract 2025-39601-44639. JF was supported by The Veterinary Scholars Research Program at The Ohio State University (NIH T35 5T35OD010977). The endemic swine IAVs were collected through funding provided by the Centers of Excellence for Influenza Research and Response (CEIRR), National Institute of Allergy and Infectious Diseases, National Institutes of Health, Department of Health and Human Services under contract 75N93021C00016. We thank the Comparative Pathology & Digital Imaging Shared Resource at The Ohio State University Comprehensive Cancer Center, Columbus, OH for the histology and immunohistochemistry microscopy, which was supported by The Ohio State University Comprehensive Cancer Center and the National Institutes of Health (P30 CA016058).

## Author contributions

Conceptualization, J.A.F., C.J.W; Data curation J.A.F., M.L., E.M.K., D.H.; Formal analysis, J.A.F., M.L., E.M.K., C.J.W.; Funding acquisition, C.J.W., A.S.B.; Investigation, J.A.F., M.L., E.M.K., D.H.; Methodology J.A.F., M.L., E.M.K., D.H., I.C.D., K.N.C.; Supervision, C.J.W., A.S.B., A.R.P.; Resources, A.R.P, I.C.D., A.S.B., C.J.W.; Visualization, J.A.F., M.L., E.M.K., K.N.C.; Writing – original draft, C.J.W, J.F., M.L., K.N.C.; Writing – review and editing, all.

## Methods

### Generation and culture of precision-cut lung slices (PCLS)

Swine lungs were collected from apparently healthy pigs immediately after harvest at a local meat processing facility and transported on ice to the laboratory. Lung lobes were rinsed thoroughly with sterile phosphate-buffered saline (PBS; Corning, #21030CV) and perfused through the major airways to remove residual blood and debris. Lobes were then trimmed into ∼2x2-inch sections using a sterile scalpel. A 2% (w/v) low-melting-point agarose solution (Invitrogen, #16520-100) was prepared in Roswell Park Memorial Institute medium (RPMI; Millipore Sigma, #R8758), heated until dissolved, and maintained at 42 °C in a bead bath to prevent solidification. Using a 3 mL syringe fitted with a 21-gauge needle, the agarose solution was slowly injected into visible airways until the tissue became firm to the touch. Inflated sections were placed on ice for 1 h to allow agarose solidification. The solidified tissue was then trimmed into smaller cubes, re-inflated with agarose, and cooled for an additional 30 min on ice. Agarose-embedded tissue blocks were placed into cylindrical molds and loaded into a Compresstome® (model VF-310-0Z, Precisionary Instruments). The slicing chamber was filled with ice-cold PBS and surrounded by ice packs to maintain low temperature. Slices of 400 μm thickness were generated using a stainless-steel blade (speed setting 2; oscillation setting 6). Individual slices were transferred into 12-well culture plates containing 1 mL of RPMI supplemented with 10% fetal bovine serum (FBS; Millipore Sigma, #F2442-500ML) and 1% Antibiotic-Antimycotic (Anti-Anti; Gibco, #15240-062). To ensure uniformity, an 8 mm biopsy punch was used to core each slice to a consistent size, and then cored slices were transferred to a new 12-well culture dish. The culture medium was replaced three times over a 3 h period (once every hour) to remove residual agarose. Tissues were then incubated overnight at 37 °C and 5% CO_2_ to allow recovery prior to experimental use.

### Cryopreservation and thawing of precision-cut lung slices

Swine precision-cut lung slices (PCLS) were cryopreserved for future use. After preparation, media refreshment, and overnight incubation as described above, individual tissue slices were transferred to cryovials containing 1.5 mL of freezing medium (RPMI supplemented with 10% FBS, 1% Anti-Anti, and 10% DMSO). Residual agarose surrounding the tissues was carefully removed prior to cryopreservation. Cryovials were placed in a controlled-rate freezing container (Mr. Frosty™) and stored at −80 °C overnight before transfer to liquid nitrogen for long-term storage. For thawing, cryovials were rapidly warmed in a 37 °C water bath, and the contents were gently transferred into a single well of a 12-well plate containing 1 mL of RPMI supplemented with 10% FBS and 1% Anti-Anti. After 1 h of incubation at 37 °C and 5% CO₂, the culture medium was replaced with fresh medium, and slices were incubated overnight to allow recovery before experimental use.

### Viruses used in this study

The human influenza A virus strains A/California/08/2009 “pdm09” and A/PR/8/34 “PR8” (non-mouse adapted) were obtained from BEI resources (cat# VR-1895 and VR-95, respectively). The endemic swine IAVs A/swine/Indiana/15TOSU1184/2015 “1184” (GenBank ID KX960572.1), A/swine/Iowa/19TOSU2799/2019 “2799” (GenBank ID MT907895.1), and A/swine/Ohio/18TOSU4522/2018 “4522” (GenBank ID MH922878.1) were collected through active surveillance. Virus stocks with unknown plaque formation unit (PFU) titers, denoted here as “passage 0” (P0), were propagated on MDCK cells as follows. Briefly, P0 virus stocks (100 μL) were diluted in 3 mL of Eagle’s minimum essential medium (EMEM, Corning, #10-009-CV) containing 1% Penicillin Streptomycin (Pen Strep, Gibco, # 15140163) and 0.3% Bovine Serum Albumin (BSA). Diluted virus stocks were then exposed to Madin Darby Canine Kidney (MDCK) cells (7.5x10^6^ cells plated in 56.7-cm2 dishes; CELLTREAT, #CT-229106) for 1 h at 37 °C with 5% CO_2_ with gentle rocking every 15 min. Following incubation, the inocula was removed and replaced with 10 mL of growth media (EMEM + 1% Pen Strep + 0.3% BSA + 1 μg/mL TPCK Trypsin [ThermoFisher Scientific, #P120233]). Cell supernatants were harvested 2-3 days later (>80% cytopathic effect) and clarified by centrifugation at 1000 x *g* for 5 min at 4 °C. This P1 stock was titrated by plaque assay and a P2 “working stock” (from which all experiments were performed) was generated by exposing MDCK cells (MOI = 0.001) as described above and passing clarified supernatant through a 0.45 μm filter. All virus stocks were maintained at -80 °C in single-use aliquots.

### Virus stock titration by plaque assay

MDCK cells were seeded at 1x10^6^ cells per well in 6-well plates. The following day, cell culture medium was removed, and cells were washed once with sterile PBS. Virus samples were 10-fold serial-diluted in serum-free EMEM, and 800 μL of diluted samples were added to each well of cells, followed by incubation at 37 °C for 1 h, with gentle rocking every 15 min. Inocula were then removed, and a volume of 3 mL of overlay medium (final concentration 1X phenol-free EMEM [Neta Scientific, #QB-115-073-101] + L-glutamine [Life Technologies, #25030081] + Pen Strep + 1.4% Avicel [IFF Pharma Solutions, #RC-581]) was added to each well, followed by undisturbed incubation at 37 °C in a humidified 5% CO_2_ atmosphere for 2 days. Following incubation, the overlay medium was removed, cells were then washed twice with PBS and fixed/stained (final concentration 20% MeOH, 0.2% crystal violet [ThermoFisher Scientific, #447570-500]) for 30 min. Plaques were enumerated following the removal of the fix/stain solution and gentle washing with DI water.

### Virus infection of precision-cut lung slices

Virus inocula were prepared by diluting each viral isolate in culture media (RPMI supplemented with 1% heat-inactivated FBS and 1% Anti-Anti) to a final concentration of 3x10^4^ forming units (PFU) in 300 μL. Before infection, the culture media was aspirated, and each slice was gently washed with 1 mL PBS. After removing the PBS, 300 μL of the viral inoculum was added per well and plates were incubated at 37°C for 1 hour with gentle rocking every 15 minutes to facilitate virus adsorption. Following incubation, inocula were aspirated, and slices were washed three times with 1 mL PBS to remove unbound virus. Fresh culture media was then added (1 mL per well) and the tissues were allowed to incubate at 37°C, 5% CO_2_. Supernatants (55 μL) were collected at 1-hour post-infection (hpi) to serve as the baseline, and at 8, 24, 72, and 96 hpi to monitor viral release kinetics. At 96 hpi, supernatants were removed, and slices were washed three times with PBS. Three replicate tissue slices per donor were collected, pooled, and homogenized for RNA extraction to quantify intracellular viral RNA. All collected media and tissue homogenates were stored at −80 °C until analysis by rRT-PCR.

### Viability assays

To determine tissue viability over time in precision-cut lung slices, lactate dehydrogenase (LDH) release was measured from mock-infected wells. Following the manufacturer’s instructions (LDH-Glo™ Cytotoxicity Assay, Promega, Cat# J2380), 2 μL of culture medium was collected from three replicate wells at 1-, 24-, 72-, and 96-hour timepoints and diluted 100-fold by adding 198 μL of LDH storage buffer to each sample. Collected samples were stored at −20 °C and batched together during assessment. On the day of the assay, samples were thawed, and 12.5 μL of each was transferred into a white, flat-bottom 384-well assay plate. An equal volume (12.5 μL) of LDH Detection Reagent was then added to each well, followed by a 60-minute incubation at room temperature. Luminescence was measured using an Infinite® 200 PRO microplate reader (Tecan) controlled by i-control™ software.

### Virus detection by rRT-PCR

Frozen samples were thawed quickly in a 37 °C dry bead bath. RNA was extracted from 50 μL of sample using the MagMAX™ Viral/Pathogen Nucleic Acid Isolation Kit (Applied Biosystems, #A48310) with the Thermo Scientific KingFisher Flex, according to the manufacturer’s protocol. The extracted RNA for each sample was evaluated for IAV RNA abundance via real-time reverse transcriptase PCR (rRT-PCR) using the VetMAX-Gold SIV Detection Kit (Life Technologies, # 4415200), according to the manufacturer’s protocol. The reactions were performed on a 7500 Fast real-time PCR system. Cycle threshold (Ct) values were set by taking 5% of the positive control at cycle 40. Samples with a Ct of ≤40 were considered positive.

### Histology and immunohistochemistry

PCLS were infected with H1N1 A/PR/8/34 “PR8” as described above. At the indicated timepoints, PCLS were washed with PBS, fixed with 10% neutral buffered formalin for 24 h, and then transitioned to 70% ethanol. Fixed PCLS samples were routinely processed on a Leica Histocore Peloris Tissue Processor, embedded in paraffin wax, sectioned by a certified histotechnologist, stained with Hematoxylin & Eosin (H&E) and coverslipped on a Leica HistoCore SPECTRA and CV Autostainer/Automated Coverslipper. CD45 immunohistochemical labeling (Abcam primary antibody product no. AB10558, Lot no. 3448663-1, 1 μg/ml working concentration) was performed by a certified histotechnologist/immunohistochemist and executed on a BioCare Medical intelliPATH FLX Autostainer platform using a quality controlled and assured automated protocol. Labeling was visualized using a horse-anti-rabbit secondary antibody (Secondary Vector Laboratories antibody product no. MP-6401, lot no. Wovus29756) and DAB chromogen with hematoxylin counterstain. All histology and IHC procedures were performed by the Ohio State University Comprehensive Cancer Center Comparative Pathology & Digital Imaging Shared Resource by certified technical staff and supervised by a board-certified comparative pathologist.

## References

1. Global Health Estimates: Life expectancy and leading causes of death and disability https://www.who.int/data/gho/data/themes/mortality-and-global-health-estimates.

2. Webby, R.J., and Uyeki, T.M. (2024). An Update on Highly Pathogenic Avian Influenza A(H5N1) Virus, Clade 2.3.4.4b. J Infect Dis 230, 533–542. 10.1093/infdis/jiae379.

3. Peacock, T.P., Moncla, L., Dudas, G., VanInsberghe, D., Sukhova, K., Lloyd-Smith, J.O., Worobey, M., Lowen, A.C., and Nelson, M.I. (2025). The global H5N1 influenza panzootic in mammals. Nature 637, 304–313. 10.1038/s41586-024-08054-z.

4. Liu, M., Faris, J.G., Panfil, A.R., and Warren, C.J. (2026). How new approach methods are reshaping virology research. J Virol 100, e0132625. 10.1128/jvi.01326-25.

5. Alsafadi, H.N., Uhl, F.E., Pineda, R.H., Bailey, K.E., Rojas, M., Wagner, D.E., and Königshoff, M. (2020). Applications and approaches for three-dimensional precision-cut lung slices. Disease modeling and drug discovery. Am J Respir Cell Mol Biol 62, 681–691. 10.1165/rcmb.2019-0276TR.

6. Vierhout, M., Ayoub, A., Ali, P., Kumaran, V., Naiel, S., Isshiki, T., Koenig, J.F.E., Kolb, M.R.J., and Ask, K. (2024). A novel ex vivo approach for investigating profibrotic macrophage polarization using murine precision-cut lung slices. Biochem Biophys Res Commun 741, 151038. 10.1016/j.bbrc.2024.151038.

7. Mansouri, S., Karger, A., Ruppert, C., Schneider, M.A., Weigert, A., Nandigama, R., Aliraj, B., Strotmann, L., Cherian, A.V., Pruefer, D., et al. (2025). Living human lung slices for ex vivo modelling of lung cancer. JCI Insight 10, e190703. 10.1172/jci.insight.190703.

8. Koziol-White, C., Gebski, E., Cao, G., and Panettieri, R.A. (2024). Precision cut lung slices: an integrated *ex vivo* model for studying lung physiology, pharmacology, disease pathogenesis and drug discovery. Respir Res 25, 231. 10.1186/s12931-024-02855-6.

9. Liu, G., Betts, C., Cunoosamy, D.M., Åberg, P.M., Hornberg, J.J., Sivars, K.B., and Cohen, T.S. (2019). Use of precision cut lung slices as a translational model for the study of lung biology. Respir Res 20, 162. 10.1186/s12931-019-1131-x.

10. Majorova, D., Atkins, E., Martineau, H., Vokral, I., Oosterhuis, D., Olinga, P., Wren, B., Cuccui, J., and Werling, D. (2021). Use of Precision-Cut Tissue Slices as a Translational Model to Study Host-Pathogen Interaction. Front Vet Sci 8, 686088. 10.3389/fvets.2021.686088.

11. Meurens, F., Summerfield, A., Nauwynck, H., Saif, L., and Gerdts, V. (2012). The pig: a model for human infectious diseases. Trends Microbiol 20, 50–57. 10.1016/j.tim.2011.11.002.

12. Judge, E.P., Hughes, J.M.L., Egan, J.J., Maguire, M., Molloy, E.L., and O’Dea, S. (2014). Anatomy and bronchoscopy of the porcine lung. A model for translational respiratory medicine. Am J Respir Cell Mol Biol 51, 334–343. 10.1165/rcmb.2013-0453TR.

13. Mena, I., Nelson, M.I., Quezada-Monroy, F., Dutta, J., Cortes-Fernández, R., Lara-Puente, J.H., Castro-Peralta, F., Cunha, L.F., Trovão, N.S., Lozano-Dubernard, B., et al. (2016). Origins of the 2009 H1N1 influenza pandemic in swine in Mexico. Elife 5, e16777. 10.7554/eLife.16777.

14. Garten, R.J., Davis, C.T., Russell, C.A., Shu, B., Lindstrom, S., Balish, A., Sessions, W.M., Xu, X., Skepner, E., Deyde, V., et al. (2009). Antigenic and genetic characteristics of swine-origin 2009 A(H1N1) influenza viruses circulating in humans. Science 325, 197–201. 10.1126/science.1176225.

15. Smith, G.J.D., Vijaykrishna, D., Bahl, J., Lycett, S.J., Worobey, M., Pybus, O.G., Ma, S.K., Cheung, C.L., Raghwani, J., Bhatt, S., et al. (2009). Origins and evolutionary genomics of the 2009 swine-origin H1N1 influenza A epidemic. Nature 459, 1122–1125. 10.1038/nature08182.

16. Cao, X., Coyle, J.P., Xiong, R., Wang, Y., Heflich, R.H., Ren, B., Gwinn, W.M., Hayden, P., and Rojanasakul, L. (2021). Invited review: human air-liquid-interface organotypic airway tissue models derived from primary tracheobronchial epithelial cells-overview and perspectives. In Vitro Cell Dev Biol Anim 57, 104–132. 10.1007/s11626-020-00517-7.

17. Crystal, R.G., Randell, S.H., Engelhardt, J.F., Voynow, J., and Sunday, M.E. (2008). Airway epithelial cells: current concepts and challenges. Proc Am Thorac Soc 5, 772–777. 10.1513/pats.200805-041HR.

18. Li, C., Huang, J., Yu, Y., Wan, Z., Chiu, M.C., Liu, X., Zhang, S., Cai, J.-P., Chu, H., Li, G., et al. (2023). Human airway and nasal organoids reveal escalating replicative fitness of SARS-CoV-2 emerging variants. Proc Natl Acad Sci U S A 120, e2300376120. 10.1073/pnas.2300376120.

19. Zhang, L., Peeples, M.E., Boucher, R.C., Collins, P.L., and Pickles, R.J. (2002). Respiratory syncytial virus infection of human airway epithelial cells is polarized, specific to ciliated cells, and without obvious cytopathology. J Virol 76, 5654–5666. 10.1128/jvi.76.11.5654-5666.2002.

20. Griggs, T.F., Bochkov, Y.A., Basnet, S., Pasic, T.R., Brockman-Schneider, R.A., Palmenberg, A.C., and Gern, J.E. (2017). Rhinovirus C targets ciliated airway epithelial cells. Respir Res 18, 84. 10.1186/s12931-017-0567-0.

21. Matrosovich, M.N., Matrosovich, T.Y., Gray, T., Roberts, N.A., and Klenk, H.-D. (2004). Human and avian influenza viruses target different cell types in cultures of human airway epithelium. Proc Natl Acad Sci U S A 101, 4620–4624. 10.1073/pnas.0308001101.

22. Sanders, C.G., Liu, M., Fusco, J.A., Ohl, E.M., Tarbuck, N.N., King, E.M., Huey, D., Fabrizio, T.P., Chen, P., Panfil, A.R., et al. (2026). Efficient replication of influenza D virus in the human airway underscores zoonotic potential. Proc Natl Acad Sci U S A 123, e2530325123. 10.1073/pnas.2530325123.

23. Alsafadi, H.N., Uhl, F.E., Pineda, R.H., Bailey, K.E., Rojas, M., Wagner, D.E., and Königshoff, M. (2020). Applications and Approaches for Three-Dimensional Precision-Cut Lung Slices. Disease Modeling and Drug Discovery. Am J Respir Cell Mol Biol 62, 681–691. 10.1165/rcmb.2019-0276TR.

24. Ma, W., Kahn, R.E., and Richt, J.A. (2008). The pig as a mixing vessel for influenza viruses: human and veterinary implications. J Mol Genet Med 3, 158–166.

25. Geiger, N., Diesendorf, V., Roll, V., König, E.-M., Obernolte, H., Sewald, K., Breidenbach, J., Pillaiyar, T., Gütschow, M., Müller, C.E., et al. (2023). Cell Type-Specific Anti-Viral Effects of Novel SARS-CoV-2 Main Protease Inhibitors. Int J Mol Sci 24, 3972. 10.3390/ijms24043972.

26. Löw, K., Möller, R., Stegmann, C., Becker, M., Rehburg, L., Obernolte, H., Schaudien, D., Oestereich, L., Braun, A., Kunz, S., et al. (2023). Luminescent reporter cells enable the identification of broad-spectrum antivirals against emerging viruses. J Med Virol 95, e29211. 10.1002/jmv.29211.

27. Geiger, N., König, E.-M., Oberwinkler, H., Roll, V., Diesendorf, V., Fähr, S., Obernolte, H., Sewald, K., Wronski, S., Steinke, M., et al. (2022). Acetylsalicylic Acid and Salicylic Acid Inhibit SARS-CoV-2 Replication in Precision-Cut Lung Slices. Vaccines (Basel) 10, 1619. 10.3390/vaccines10101619.

28. Chen, P., K C, M, and Peeples, M.E. (2025). Well-Differentiated Primary Human Airway Epithelial Cultures for Respiratory Syncytial Virus (RSV) Infection Studies. Methods Mol Biol 2948, 249–265. 10.1007/978-1-0716-4666-3_16.

29. Chang, S.-Y., Chang, W.-H., Yang, D.C., Hong, Q.-S., Hsu, S.-W., Wu, R., and Chen, C.-H. (2025). Autologous precision-cut lung slice co-culture models for studying macrophage-driven fibrosis. Front Physiol 16, 1526787. 10.3389/fphys.2025.1526787.

